# The Type VI Secretion System of the Emerging Pathogen *Stenotrophomonas maltophilia* has Antibacterial Properties

**DOI:** 10.1101/2023.05.30.542968

**Authors:** Cristian V. Crisan, Daria Van Tyne, Joanna B. Goldberg

## Abstract

Antagonistic behaviors between bacterial cells can have profound effects on microbial populations and disease outcomes. Polymicrobial interactions may be mediated by contact-dependent proteins with antibacterial properties. The Type VI Secretion System (T6SS) is a macromolecular weapon used by Gram-negative bacteria to translocate proteins into adjacent cells. The T6SS is used by pathogens to escape immune cells, eliminate commensal bacteria, and facilitate infection. *Stenotrophomonas maltophilia* is a Gram-negative opportunistic pathogen that causes a wide range of infections in immunocompromised patients and infects the lungs of patients with cystic fibrosis. Infections with the bacterium can be deadly and are challenging to treat because many isolates are multidrug-resistant. We found that globally dispersed *S. maltophilia* clinical and environmental strains possess T6SS genes. We demonstrate that the T6SS of an *S. maltophilia* patient isolate is active and can eliminate other bacteria. Furthermore, we provide evidence that the T6SS contributes to the competitive fitness of *S. maltophilia* against a co-infecting *Pseudomonas aeruginosa* isolate, and that the T6SS alters the cellular organization of *S. maltophilia* and *P. aeruginosa* co-cultures. This study expands our knowledge of the mechanisms employed by *S. maltophilia* to secrete antibacterial proteins and compete against other bacteria.

**IMPORTANCE:** Infections with the opportunistic pathogen *Stenotrophomonas maltophilia* can be fatal for immunocompromised patients. The mechanisms used by the bacterium to compete against other prokaryotes are not well understood. We found that the T6SS allows *S. maltophilia* to eliminate other bacteria and contributes to the competitive fitness against a co-infecting isolate. The presence of T6SS genes in isolates across the globe highlights the importance of this apparatus as a weapon in the antibacterial arsenal of *S. maltophilia*. The T6SS may confer survival advantages to *S. maltophilia* isolates in polymicrobial communities in both environmental settings and during infections.

## INTRODUCTION

*Stenotrophomonas maltophilia* is an emerging, globally dispersed opportunistic pathogen that causes infections of the lungs, brain, gastrointestinal tract, nervous system, eyes, blood, skin and bones (1–3). Bacteremia caused by the bacterium can have mortality rates as high as 65% (4). Infections are difficult to treat because many isolates are intrinsically resistant to multiple antibiotics, including cephalosporins (5–7). In some hospitals, *S. maltophilia* is the most common Gram-negative, carbapenem-resistant bacterium in patients with bacteremia (8). In 2018, the World Health Organization listed *S. maltophilia* as an important pathogen for which novel antibiotic treatments are urgently needed (6, 7).

Immunocompromised and cancer patients are especially susceptible to *S. maltophilia* infections (4, 9). Cystic fibrosis (CF), a genetic disease that affects more than 150,000 people worldwide, leads to chronic pulmonary bacterial infections (10). Approximately 10-30% of CF patients harbor *S. maltophilia* in their lungs at least once during their life (11, 12). CF pulmonary *S. maltophilia* infections can be associated with higher mortality, more severe exacerbations, and an increased probability of requiring lung transplants (13). The bacterium is also detected in sputum obtained from COVID-19 patients, and *S. maltophilia* isolates have the highest rates of multidrug resistance among bacterial species that infect COVID-19 patients (14–16).

*S. maltophilia* is ubiquitously found in many water and soil environments, and has been isolated from hospital surfaces and medical devices (17, 18). The species exhibits extensive genomic diversity and has been subdivided into 23 monophyletic lineages (19). Due to its diversity, the phylogenetic classification of the species is problematic and Gröschel et al. proposed the term “*S. maltophilia* complex” for isolates identified as *S. maltophilia* by diagnostic procedures (19). *S. maltophilia* is frequently isolated from patients with co-infecting pathogens like *Pseudomonas aeruginosa,* and both cooperative and antagonistic interactions have been observed between the two bacterial species (20–24). The *S. maltophilia* K279a blood isolate possesses a Type IV Secretion System (T4SS) with antibacterial properties (20, 21, 24). The T4SS contributes to the ability of K279a to eliminate other bacteria, including clinical *P. aeruginosa* isolates (20, 21, 24). VirB10 (part of the outer membrane pore complex) and VirD4 (an ATPase) proteins are required for the antibacterial activity of the T4SS in *S. maltophilia* K279a (20, 21, 24).

The Type VI Secretion System (T6SS) is an important proteinaceous macromolecular apparatus used by many Gram-negative pathogens to deliver toxic proteins (effectors) into target cells (25–27). A membrane complex (which includes the essential TssM protein) and a baseplate structure assemble within the membrane of cells that possess an active T6SS (28, 29). The inner tube of stacked Hcp (hemolysin-coregulated protein) hexamers is surrounded by a contractile outer sheath formed by TssB and TssC proteins (30–32). Previous studies used TssC protein sequences to determine T6SS phylogenetic relationships (33). The Secret6 database classifies T6SS sequences into four types (i, ii, iii, and iv), while type i is further divided into six subtypes (1, i2, i3, i4a, i4b and i5) (34). Contractions of the outer sheath facilitate excretion of the Hcp tube, which is capped by a VgrG (valine-glycine repeat protein) and PAAR (proline-alanine-alanine-arginine) complex (35–37). Secreted T6SS toxic proteins (effectors) interact with VgrG or Hcp proteins and can exhibit both antibacterial and anti-eukaryotic properties (25, 38, 39). Rhs (rearrangement hotspot) toxins are large, polymorphic proteins that may associate with the T6SS (40, 41). Antibacterial effectors exert their toxicity on the cell envelope (where they can damage the peptidoglycan layer, degrade lipid membranes, or form pores) and the cytoplasm (where they can alter nucleotides or interfere with protein synthesis) of competitor bacteria (39, 42–46).

In this study we performed bioinformatic searches using NCBI databases and identified *S. maltophilia* isolates from both patient and environmental sources that possess T6SS genes. We found that a clinical *S. maltophilia* complex strain utilizes a T6SS that is active under standard laboratory conditions and is employed to eliminate other bacteria like *Escherichia coli* and *Burkholderia cenocepacia*. We also observed that the T6SS confers a competitive advantage to *S. maltophilia* against a co-infecting *P. aeruginosa* strain obtained from the same patient. We used confocal microscopy to determine that the T6SS alters the spatial organization of co-cultures containing both *S. maltophilia* and *P. aeruginosa*. Although T6SS genes have been previously reported in *S. maltophilia* complex strains (47, 48), experimental studies to demonstrate their function are lacking. Here, we provide the first evidence for the activity of the T6SS in the *Stenotrophomonas* genus.

## RESULTS

### T6SS genes are found in globally distributed *S. maltophilia* patient and environmental isolates

To analyze the distribution of T6SS genes in *S. maltophilia* complex isolates, we used the sequence of the TssC sheath protein from *Xanthomonas citri* to search for homologous proteins encoded by approximately 1000 *S. maltophilia* complex genomes from RefSeq and Genbank databases (49, 50)*. X. citri* belongs to the same *Xanthomonadaceae* family as *S. maltophilia* and possesses a T6SS that contributes to resistance against eukaryotic predators (51). We discovered that 61 *S. maltophilia* complex isolates from animal, human, plant, and environmental sources contain one or two copies of TssC (Fig. 1A, Supp. Table 1). Strains harboring T6SS genes are dispersed across multiple countries from Africa, Asia, Australia, Europe, and America and some were obtained from patient lung, blood, urine, and wound samples (Fig. 1B and Supp. Table 1). Three CF isolates (B4, B5, and H_59_creteil) possess T6SS genes (Fig. 1A and Supp. Table 1).

**Figure 1.**
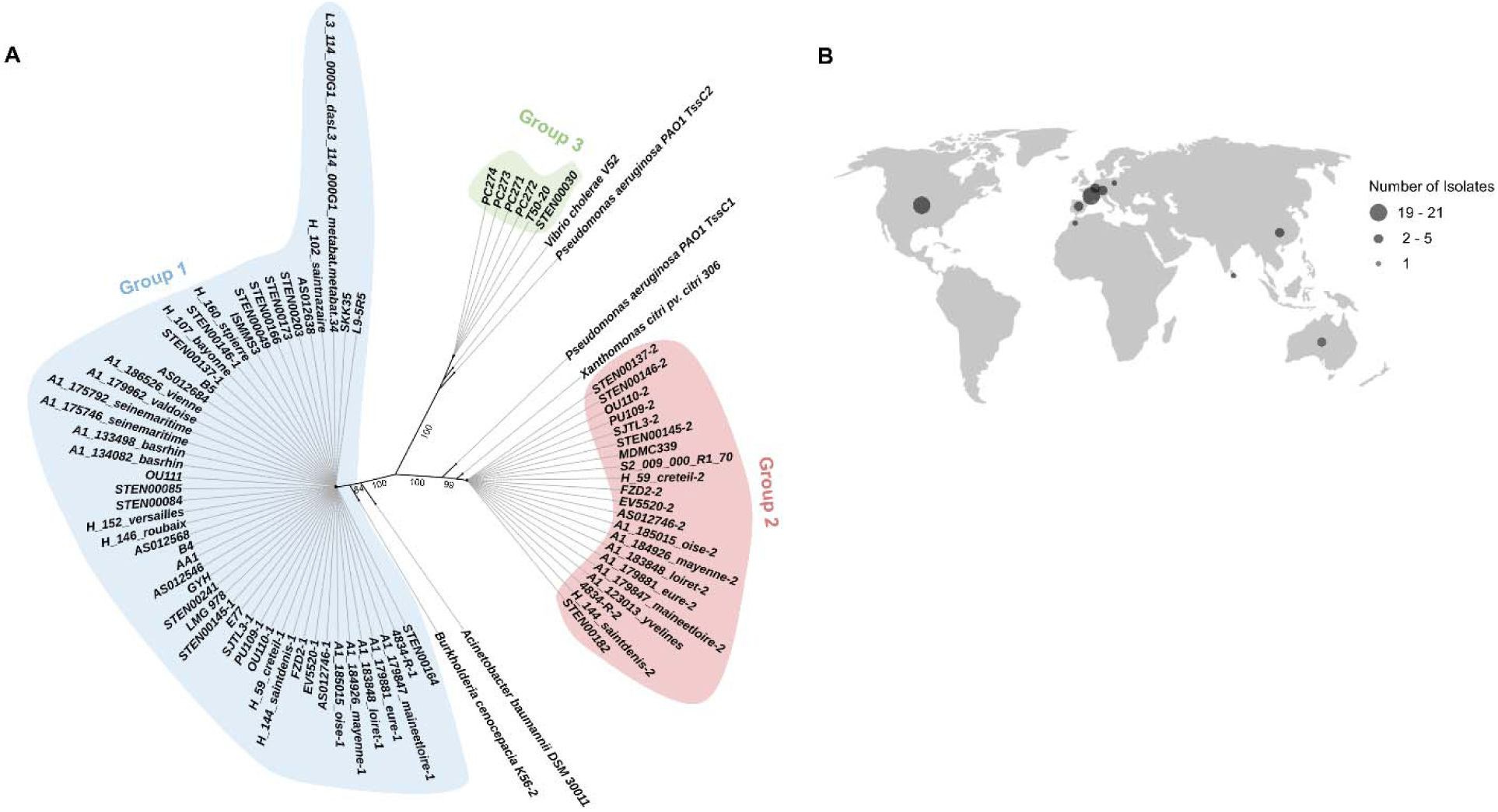
Patient and environmental *S. maltophilia* complex isolates from diverse geographic locations harbor T6SS genes. A) A BlastP search was conducted to identify *S. maltophilia* complex TssC proteins. TssC sequences of the indicated strains were aligned, and a phylogenetic tree was constructed. Branch titles designate the strain name that encodes the respective TssC homologue. Branch numbers indicate support values. Strains that harbor a TssC from groups 1 and 2 are indicated by a “-1” or “-2”, respectively, after the strain name. B) The number of *S. maltophilia* complex isolates with T6SS genes from each country is displayed.

Based on their amino acid sequence, *S. maltophilia* complex TssC proteins cluster into three distinct phylogenetic groups: group 1, group 2, or group 3 (Fig. 1A). Group 1 TssC proteins are the most common and share homology to the TssC of *Burkholderia cenocepacia* and *Acinetobacter baumannii*, whose T6SSs belong to the i4b subtype (34, 52, 53). By contrast, TssC proteins from group 2 cluster with TssC1 from *P. aeruginosa*, which belongs to the i3 T6SS subtype, and TssC from *X. citri* (26, 51). Four *S. maltophilia* isolates harbor only a single TssC from group 2 and no TssC from group 1 (Fig. 1, Supp. Table 2). Finally, *S. maltophilia* complex TssC proteins from group 3 are clustered with TssC2 from *P. aeruginosa* and TssC from *V. cholerae*, which are part of the i1 T6SS subtype (34).

We surveyed across *S. maltophilia* complex strains that encode TssC proteins for the presence of T4SS (VirB10 and VirD4) proteins. All *S. maltophilia* complex isolates with a TssC from group 3, as well as isolates MDMC339 (with a TssC from group 2) and AS012546 (with a TssC from group 1), encode T4SS proteins (Fig. 2). We next performed an average nucleotide identity (ANI) analysis using *S. maltophilia* complex genomes with T6SS genes and included the genome of K279a (which possesses T4SS but not T6SS genes) (Fig. 2). Strains that encode both T6SS and T4SS proteins belong to distinct isolates that cluster near K279a, while most isolates harboring TssC proteins from groups 1 and 2 form a separate group (Fig. 2). In conclusion, *S. maltophilia* complex isolates from both clinical and environmental sources harbor diverse T6SS loci.

**Figure 2.**
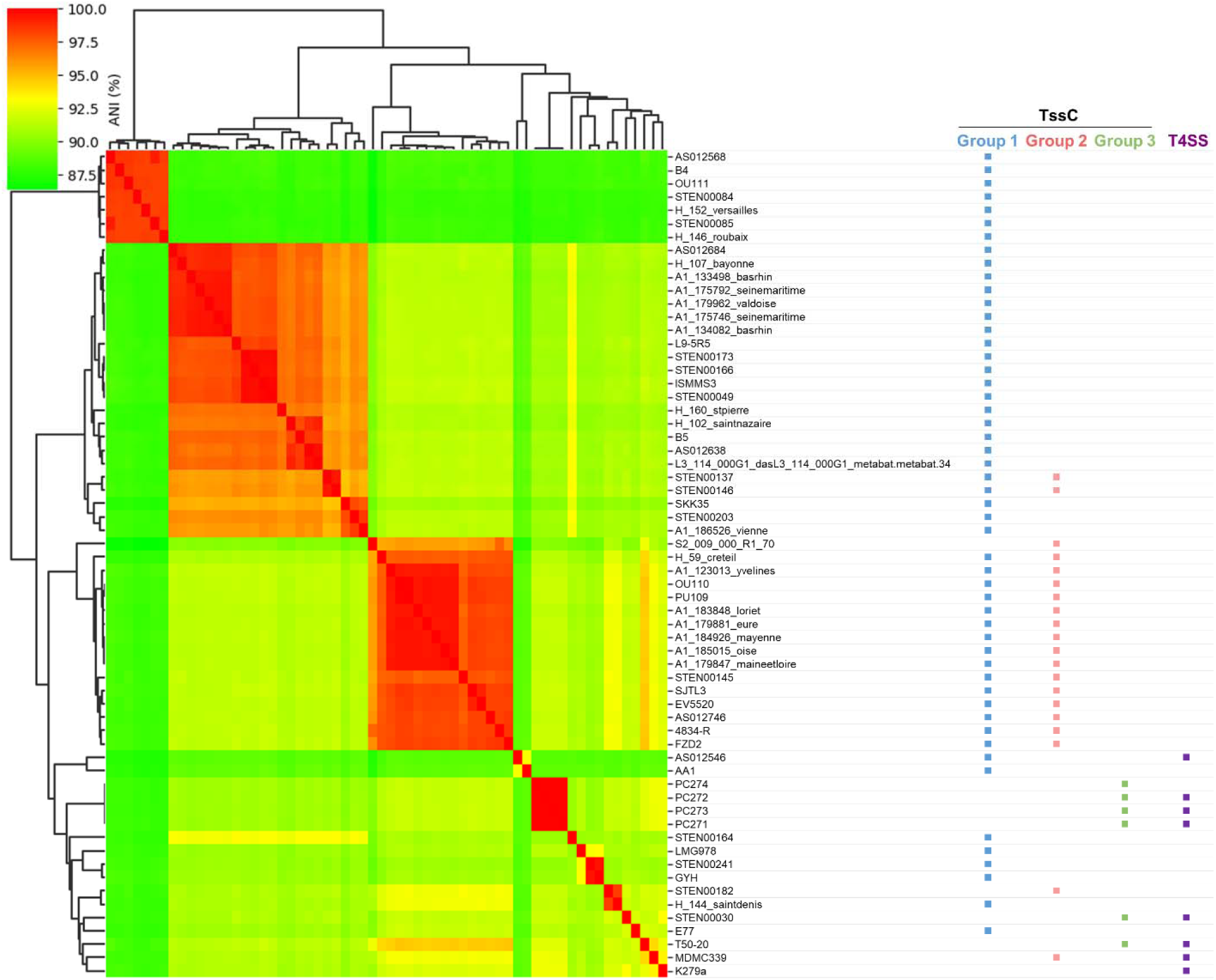
Average nucleotide identity (ANI) between 61 *S. maltophilia* complex strains with T6SS genes and *S. maltophilia* K279a. Genomic nucleotide sequences of the indicated *S. maltophilia* complex isolates were compared using FastANI and a matrix was created from the obtained values. Strains with VirB10 and VirD4 T4SS protein sequences (which are essential for the antibacterial activity of the T4SS) with >50% similarity to their homologs from K279a were considered to have a T4SS.

### STEN00241 possesses essential T6SS genes and a repertoire of predicted toxins

To further understand the organization and function of T6SS genes in *S. maltophilia* complex, we analyzed sputum isolate STEN00241, which harbors a single TssC from group 1. STEN00241 possesses a main operon with T6SS genes (referred to as T6SS-1) predicted to encode proteins that form the membrane complex, baseplate, inner tube, sheath, and tip of the apparatus (Fig. 3A). Additionally, we discovered eight alleles encoding VgrG proteins that were distributed throughout the genome (Fig. 3A and 3B). *vgrG-1* is found within the main T6SS operon, while *vgrG-2* and *vgrG-3* are located immediately downstream of the main T6SS operon (Fig. 3A). Both *vgrG-2* and *vgrG-3* are found near genes predicted to encode chaperones (DUF4123), phospholipases (DUF2235), and immunity proteins (DUF3304) (42, 54, 55). *vgrG*-4 is found immediately upstream of the T6SS main operon near a gene predicted to encode a protein with lysozyme-like function.

**Figure 3.**
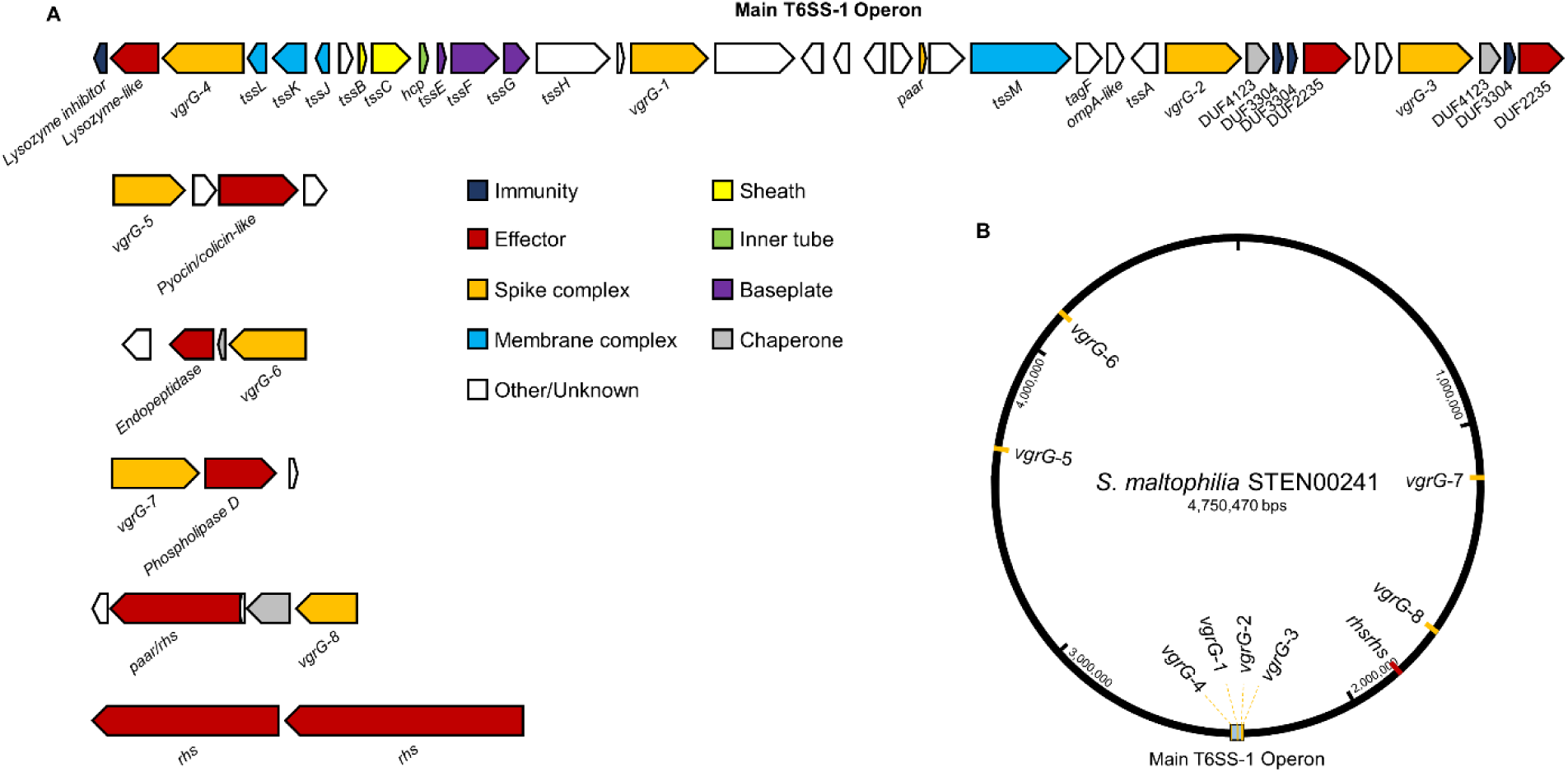
Genes encoding T6SS structural components and toxins are found in *S. maltophilia* complex STEN00241. A) The T6SS-1 cluster of STEN00241 contains T6SS genes predicted to encode essential components of the membrane complex (blue), baseplate (purple), inner tube (green), outer sheath (yellow), and tip complex (orange). Orphan *vgrG* genes are found near putative toxin and immunity genes. B) Genomic locations of the main T6SS and VgrG clusters on the genome of STEN00241.

Unlike *vgrG*1-4, other *vgrG* genes are found at locations distal to the main T6SS operon (Fig. 3B). Downstream of *vgrg-5,* a gene encoding a colicin/pyocin-like protein is found, while a putative endopeptidase is predicted to be encoded downstream of *vgrG-6.* Similarly, a gene predicted to encode a phospholipase is located downstream of *vgrG-*7. Three *rhs* genes are also found in the genome of STEN00241: one is located downstream of *vgrG-8,* while the other two are found near each other but distant from a *vgrG* gene. These findings demonstrate that *S. maltophilia* complex STEN00241 harbors essential T6SS genes as well as a diverse array of putative effectors.

### The *S. maltophilia* complex T6SS is active and displays antibacterial properties

Since we observed multiple toxins with putative antibacterial properties in the genome of STEN00241, we hypothesized that the T6SS is used to eliminate competitor bacteria. We engineered a T6SS deficient mutant of STEN00241 by deleting the *tssM* gene (Δ*tssM*), which is essential for the function of T6SS in other bacteria (52, 56). The Δ*tssM* strain has a similar growth rate to the wild type (WT) strain in liquid LB medium (Supp. Fig. 1). We co-cultured WT or Δ*tssM* STEN00241 mutant strains with *E. coli* cells and observed that the WT strain robustly eliminates *E. coli* at 37°C (Fig. 4A) and 25°C (Supp. Fig. 2). By contrast, the Δ*tssM* mutant is significantly impaired at killing *E. coli* (Fig. 4A, Supp. Fig. 2). Approximately the same number of WT and Δ*tssM* mutant STEN00241 cells are recovered following co-culture with *E. coli* (Fig. 4B). Secretion of the Hcp protein in the supernatant has been previously used to demonstrate active T6SS in other bacterial species (52, 57). We observed that WT *S. maltophilia* secretes Hcp in the supernatant, while the Δ*tssM* mutant does not (Fig. 4C).

**Figure 4.**
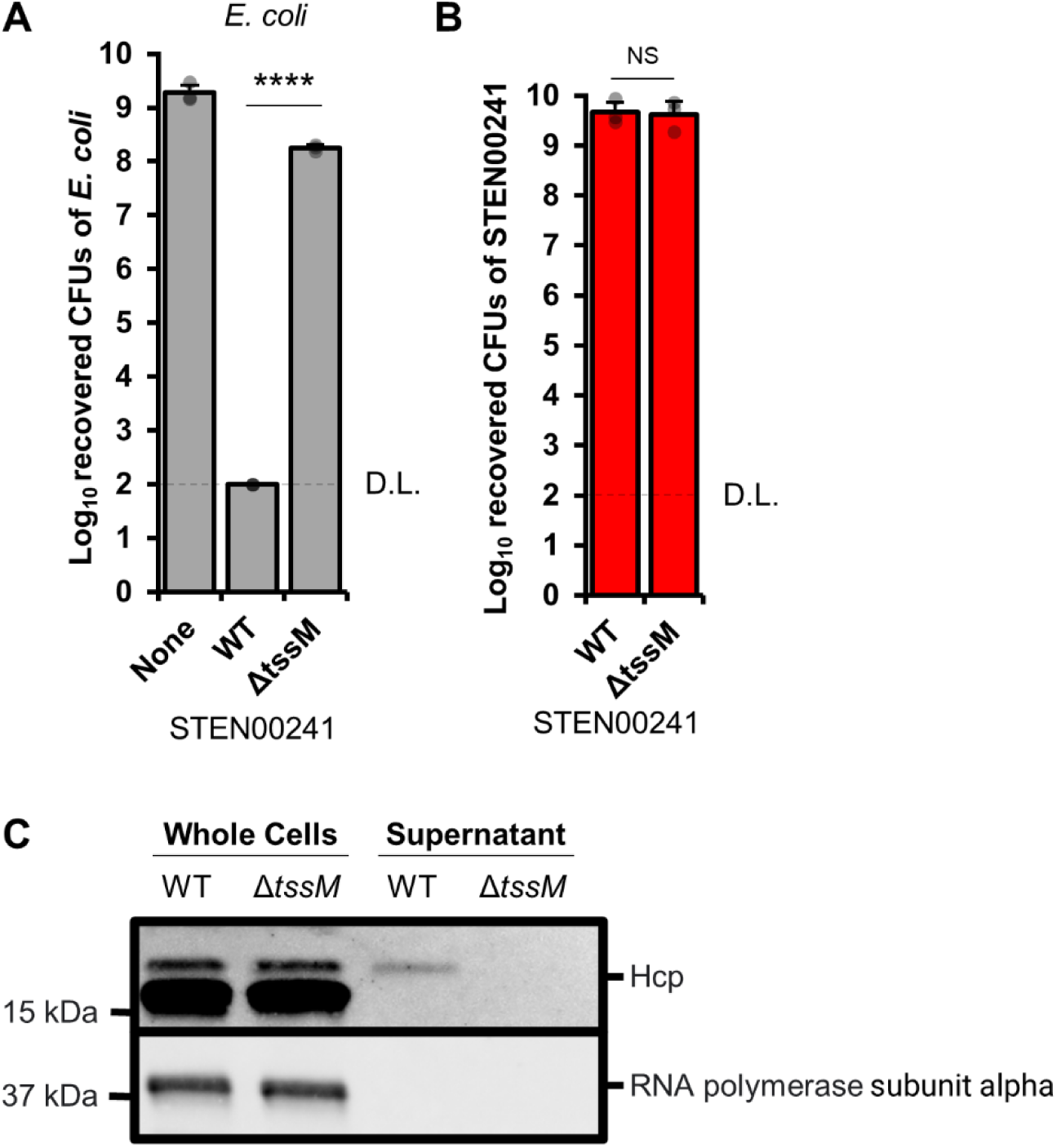
The *S. maltophilia* TssM membrane complex protein is required for eliminating heterologous bacteria and secreting Hcp. A) *E. coli* resistant to tetracycline were grown alone or in the presence of *S. maltophilia* (WT or Δ*tssM*) on solid LB medium at a 1:10 (*E. coli*:*S. maltophilia*) ratio for 20 hours at 37°C. The number of surviving *E. coli* cells was determined by plating mixtures on tetracycline plates. B) Same as in A, but the number of surviving *S. maltophilia* cells was determined by plating mixtures on imipenem plates. Three independent biological replicates were performed. For A, a one-way ANOVA with post-hoc Tukey HSD was used to determine statistical significance. For B, a Welch’s unequal variances t-test was used to determine statistical significance. **** p < 0.0001, NS – not significant (p > 0.05). D.L. – detection limit. C) Whole cells and supernatants from STEN00241 WT and Δ*tssM* were probed with antibodies against Hcp and RNA polymerase subunit alpha (cell lysis control).

STEN00241 also eliminates *B. cenocepacia* strain K56-2 in a T6SS-dependent manner (Fig. 5A). By contrast, the T6SS does not contribute to killing of the *P. aeruginosa* PA14 laboratory strain or a *P. aeruginosa* CF isolate (PA32) (Fig. 5B and 5C). Similarly, the T6SS does not affect the ability of *S. maltophilia* to eliminate the *Staphylococcus aureus* JE2 laboratory strain (Fig. 5D). These results demonstrate that STEN00241 can utilize the T6SS to eliminate some heterologous bacterial species.

**Figure 5.**
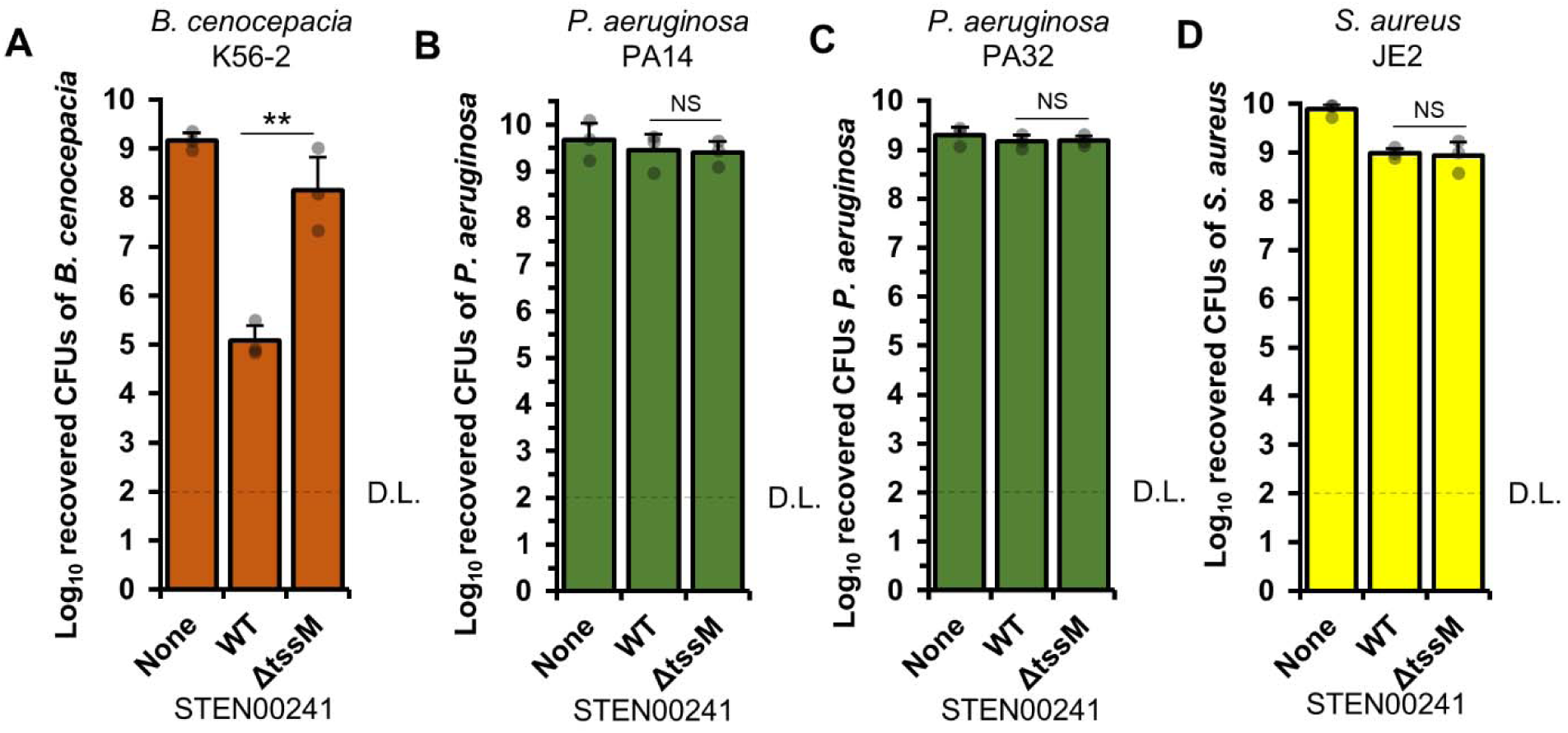
The *S. maltophilia* complex T6SS contributes to the elimination of *B. cenocepacia,* but not *P. aeruginosa* PA14, PA32, or *S. aureus*. A) *B. cenocepacia* K56-2 was grown alone or in the presence of *S. maltophilia* complex STEN00241 (WT or Δ*tssM*) at a 1:10 (*B. cenocepacia*:*S. maltophilia*) ratio on solid LB medium and incubated for 20 hours at 37°C. The number of surviving *B. cenocepacia* cells was determined by plating mixtures on Gentamicin plates. B) Same as in A), but *P. aeruginosa* PA14 used instead of *B. cenocepacia.* The number of surviving *B. cenocepacia* cells was determined by plating mixtures on chloramphenicol plates. C) Same as in B), but *P. aeruginosa* PA32 was used instead of PA14. D) Same as in A), but *S. aureus* was used instead of *B. cenocepacia*. The number of surviving *S. aureus* cells was determined by plating mixtures on *Staphylococcus* isolation agar. Three independent biological replicates were performed. A one-way ANOVA with post-hoc Tukey HSD was used to determine statistical significance. ** p < 0.01, NS – not significant (p > 0.05). D.L. – detection limit.

### The T6SS contributes to the competitive fitness of STEN00241 against a co-infecting *P. aeruginosa* isolate

Interactions that occur between two bacterial species co-isolated from the same patient can have distinct outcomes compared to interactions that occur between strains obtained from different sources (58, 59). STEN00241 was co-isolated from the same patient as *P. aeruginosa* strain PSA01136 (60). We competed STEN00241 and PSA01136 with one another and observed their dynamics. Approximately 10-fold fewer PSA01136 cells are recovered when the strain is competed against the WT STEN00241 compared to the Δ*tssM* mutant (Fig. 6A). By contrast, the number of recovered PSA01136 cells are not significantly affected by the T6SS when co-cultures are performed in liquid conditions, which allow only minimal contact to occur between cells (Supp. Fig. 3). Furthermore, we also observed that the number of recovered STEN00241 Δ*tssM* cells is significantly lower compared to the number of recovered WT *S. maltophilia* cells when co-cultured on solid medium with PSA01136, suggesting that the T6SS contributes to the ability of STEN00241 to survive against the co-infecting *P. aeruginosa* isolate (Fig. 6B).

**Figure 6.**
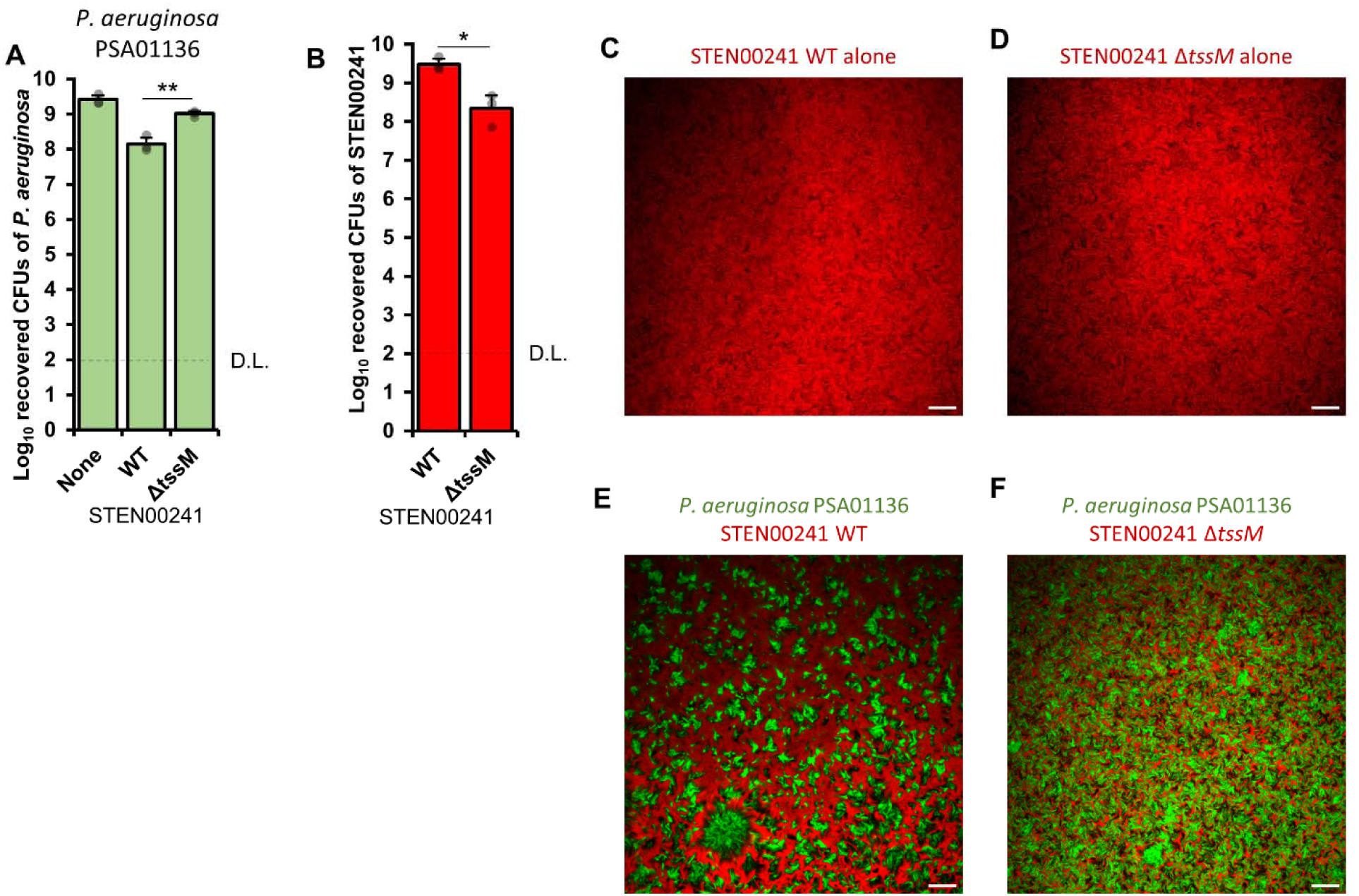
*S. maltophilia* STEN00241 uses the T6SS to compete against a *P. aeruginosa* co-infecting isolate. A) *P. aeruginosa* PSA01136 was grown alone or in the presence of *S. maltophilia* complex STEN00241 (WT or Δ*tssM*) at a 1:10 (*P. aeruginosa*:*S. maltophilia*) ratio on solid LB medium and incubated for 20 hours at 37°C. The number of surviving *P. aeruginosa* PSA01136 cells was determined by plating mixtures on chloramphenicol plates. B) Same as above, but the number of surviving *S. maltophilia* cells was determined by plating mixtures on imipenem plates. Three independent biological replicates were performed. For A, a one-way ANOVA with post-hoc Tukey HSD was used to determine statistical significance. For B, a Welch’s unequal variances t-test was used to determine statistical significance. ** p < 0.01, * p < 0.05, NS – not significant (p > 0.05). D.L. – detection limit. For C-F, STEN00241 WT alone expressing mCherry (C), STEN00241 Δ*tssM* alone expressing mCherry (D), co-cultures between *P. aeruginosa* PSA01136 expressing GFP and STEN00241 WT expressing mCherry (E), or co-cultures between *P. aeruginosa* PSA01136 expressing GFP and *S. maltophilia* Δ*tssM* expressing mCherry (F) were visualized using a Zeiss LSM 710 upright microscope. Images were analyzed in FIJI and are representative of three independent biological replicates. Scale bar represents 100 µm.

We used confocal microscopy to observe co-cultures between STEN00241 WT and Δ*tssM* expressing mCherry and PSA01136 expressing green fluorescent protein. When PSA01136 is co-cultured with WT STEN00241, *P. aeruginosa* forms large, distinct clusters from which STEN00241 cells are mostly excluded (Fig. 6D). By contrast, when PSA01136 is co-cultured with STEN00241 Δ*tssM*, the two strains form a mixed, interwoven pattern (Fig. 6E). Taken together, these results provide evidence that the T6SS contributes to the competitive fitness of STEN00241 when co-cultured with a co-infecting *P. aeruginosa* isolate and alters interactions between the two bacteria.

### T4SS and T6SS genes are found at distinct genomic locations in *S. maltophilia* complex strains

Previous work showed that *S. maltophilia* K279a uses a T4SS to eliminate bacteria (20, 21, 24) and here we provide evidence that the T6SS of *S. maltophilia* can also display antibacterial properties. We wondered whether the two systems are found at the same genomic locations in different strains. We analyzed the complete genomes of four *S. maltophilia* isolates to compare the genomic locations of TssC genes (from groups 1-3) and T4SS gene clusters. For this comparison we used strain STEN00241, which harbors a TssC from group 1 (within the T6SS-1 cluster), strain K279a, which possesses a T4SS cluster but no T6SS genes, strain SJTL3, which has both a TssC from group 1 (within a T6SS-1 cluster) and a TssC from group 2 (within a T6SS-2 cluster), and strain T50-20, which contains a TssC from group 3 (within a T6SS-3 cluster) and a T4SS (Fig. 7).

**Figure 7.**
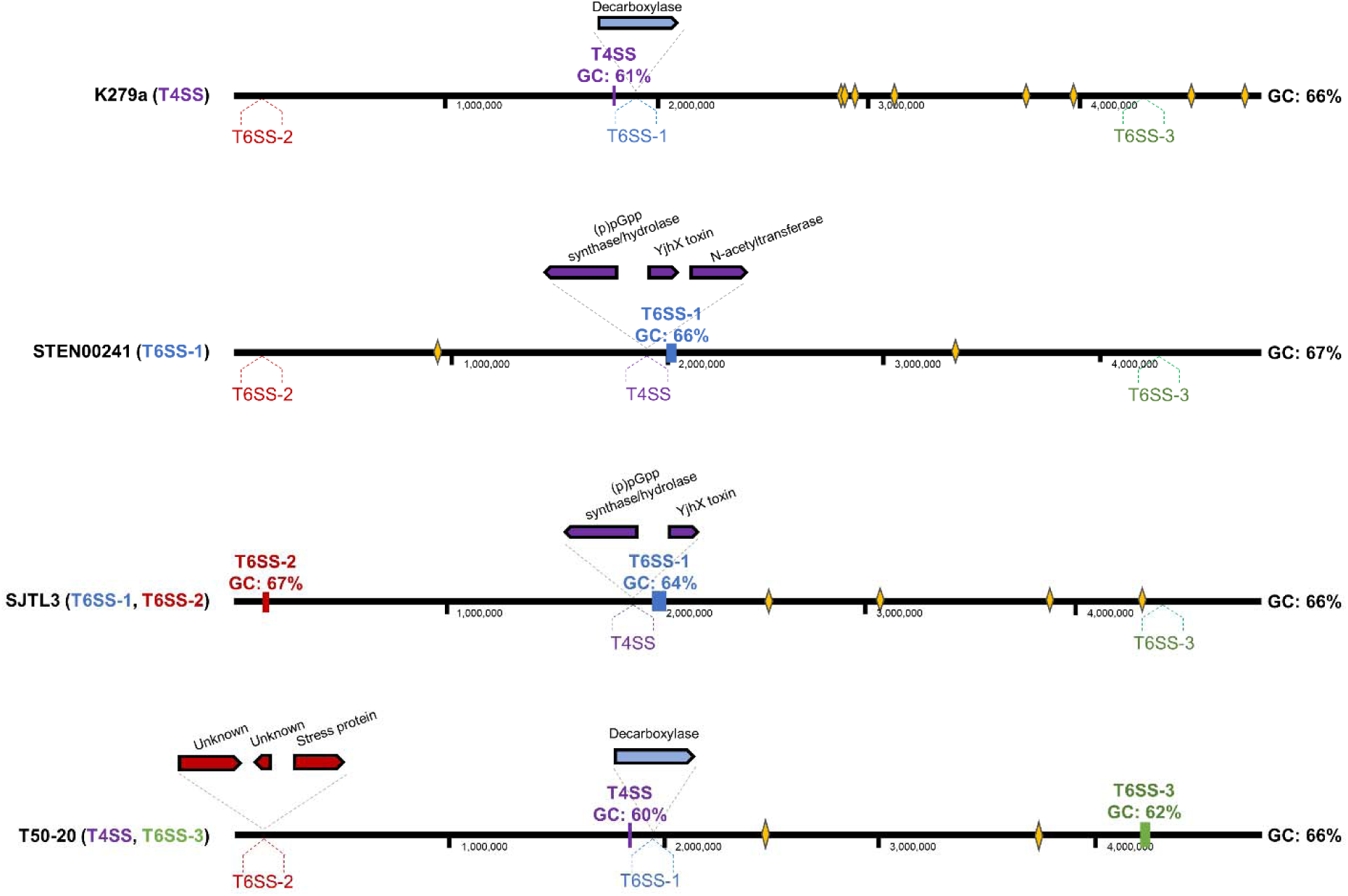
Genes coding for T4SS and T6SS components are found at conserved locations in *S. maltophilia*. T4SS (purple), T6SS-1 (blue), T6SS-2 (red), and T6SS-3 (green) genomic sequences were mapped on the genomes of *S. maltophilia* K279a, STEN00241, SJTL3, and T50-20. When sequences were missing, upstream and downstream genomic regions were mapped instead. Dashed lines and names below the solid black lines indicate the putative locations of elements that are missing from the genomes. Rectangles on the black line, gene representations, and names above the solid black line represent elements present in the genomes. Numbers below the lines correspond to genomic locations in each genome. GC percentages for each gene cluster are displayed below their respective names and whole genome GC percentages are displayed to the right of each genome. Yellow diamonds represent putative phage elements predicted by PHASTER (61).

*S. maltophilia* isolates that lack T6SS or T4SS elements possess upstream and downstream regions of the missing sequences at approximately the same genomic locations as strains that harbor T6SS or T4SS genes (Fig. 7). SJTL3 and STEN00241 appear to have exchanged the T4SS cluster for genes encoding a (p)pGpp synthase/hydrolase and a toxin from a toxin-antitoxin (TA) system (Fig. 7). Strain T50-20 encodes a stress protein in the place of a T6SS-2 module, while both T50-20 and K279a encode a decarboxylase in place of the T6SS-1 operon. The GC content of T4SS and T6SS modules are similar to that of the entire genomes, except for the GC content of the T4SS and T6SS-3 clusters, which have 60-62% GC compared to the 66% genomic GC content of K279a and T50-20. We did not detect phage-related sequences in the immediate vicinity (∼50,000 nucleotides) of T4SS or T6SS modules (Fig. 7, yellow diamonds), suggesting that these elements were not acquired by transduction events.

## DISCUSSION

The study presented here describes the T6SS as a novel antibacterial weapon in the arsenal of *S. maltophilia* complex. In both host and environmental settings, bacteria like *S. maltophilia* live in polymicrobial communities where interactions with other cells influence the ability of a species to persist (62). These interactions are often antagonistic and may be mediated by diffusible antibacterial small molecules or by contact-dependent proteins that are transferred between cells to intoxicate competitors (25, 62). Prior studies have identified T6SS genes in other *S. maltophilia* complex strains, but it is unclear what role the T6SS plays in those isolates (47, 48).

We report here that T6SS genes are found in geographically diverse *S. maltophilia* complex isolates obtained from both patient and non-patient sources, suggesting that the T6SS might confer a competitive advantage against other bacteria during infection and during survival in the environment. This hypothesis is supported by our finding that the T6SS is important for the elimination of *E. coli* at both 25°C and 37°C. *P. aeruginosa* and *B. cenocepacia* also possess active T6SS clusters that may contribute to pathogenicity and allow those pathogens to outcompete other bacteria (25, 27). *P. aeruginosa* Hcp proteins and antibodies against Hcp have been detected in the sputum of CF patients (26). *P. aeruginosa* CF isolates with loss-of-function mutations in T6SS regulator genes become susceptible to killing by species from the *Burkholderia cepacia* complex (27). It is currently unclear if the T6SS of *S. maltophilia* is important in infections such as those seen in CF patients.

*P. aeruginosa* and *S. maltophilia* complex species are frequently isolated from the same patients and both cooperative and antagonistic interactions between the two species have been described (20–22). During mouse lung infections, *P. aeruginosa* can increase *S. maltophilia* proliferation and associated immune responses (22). Patient isolates of *P. aeruginosa* and *S. maltophilia* have been observed to share genomic sequences, suggesting that inter-species horizontal gene transfer is common among these organisms (63). However, competitive interactions can also occur between these two pathogens. *S. maltophilia* K279a uses the T4SS to deliver toxins with predicted lipase and lysozyme-like activity to intoxicate and kill *P. aeruginosa* strains (20, 21, 24). We provide evidence that an *S. maltophilia* complex pulmonary isolate can utilize the T6SS to eliminate heterologous bacteria *B. cenocepacia* and to compete against a co-infecting *P. aeruginosa* isolate.

Even though the T6SS contributes to the competitive fitness of *S. maltophilia* against the co-infecting *P. aeruginosa* PSA01136 strain, it does not play a significant role in competition against the lab strain PA14 or the CF isolate PA32. Perault et al. observed that *P. aeruginosa* isolates from adult patients harbor mutations in the GacS, GacA, RetS, Fha1, and PppA proteins that regulate T6SS gene expression in *P. aeruginosa* (27). We also observed mutations in genes encoding those regulators in *P. aeruginosa* PSA01136 compared to PA14. However, the impact played by those mutations in mediating T6SS activity in PSA01136 is unclear. *P. aeruginosa* employs defense mechanisms against T6SS attacks, such as the Arc immunity pathway and stress response systems (63, 64). We suspect that variable expression of defensive and offensive systems in the co-infecting PSA01136 isolate could explain the differences we observed in their competitive fitness against STEN00241 compared to PA14. Although some exceptions have been recently identified, most T6SS are not effective at eliminating Gram-positive bacteria (65). While competition with *S. maltophilia* reduced the number of recovered *S. aureus* during co-cultures compared to monocultures, the T6SS did not affect survival of *S. aureus*.

The distribution and organization of T6SS genes across bacterial genomes is diverse. Species from the *Burkholderia* genus may harbor up to six distinct T6SS clusters, while *P. aeruginosa* strains generally possess three T6SS clusters (26, 66). *Vibrio cholerae* strains employ a single T6SS operon that encodes structural and regulatory proteins, as well as orphan *vgrG* gene loci that contain effectors with diverse functions (67). We found that *S. maltophilia* strains can encode TssC proteins from groups 1 and 2 but all isolates that have a TssC from the group 3 only have a single TssC. All *S. maltophilia* complex strains with a TssC from group 3 also harbor T4SS genes homologous to the ones used by K279a to eliminate bacteria (20, 21, 24). In other bacteria that harbor multiple T6SS clusters, each system can play a role in mediating virulence, obtaining nutrients, and conferring competitive advantages against other bacteria (68–70). It is unclear why some *S. maltophilia* complex isolates harbor T4SS genes while others possess T6SS components. Based on the four complete genomes analyzed in Fig. 7, T4SS or T6SS modules appear at conserved but distinct genomic locations, suggesting that they are not part of interchanged mobile elements. We propose that *S. maltophilia* complex strains have separate genomic locations which serve as an armory that accommodates different molecular weapons. We speculate that evolutionary pressures dictated by the environment, like competition with eukaryotic or prokaryotic cells, the energy cost of assembling and firing a T4SS or T6SS in different conditions, and the efficacy of secreted toxins might play key roles in determining which secretion system is acquired and maintained by a *S. maltophilia* complex strain. Future work will determine whether different *S. maltophilia* T6SS are required for different processes.

Results presented here enhance our understanding of the secretion systems and antibacterial weapons employed by an important emerging pathogen. We propose that strains harboring T6SS genes have competitive fitness advantages during survival in both infections and on hospital or environmental surfaces. Additional studies will elucidate how the *S. maltophilia* complex T6SS is regulated, which components of the apparatus are required for activity, which toxins are important in mediating the observed competitive fitness advantages, and how the system might affect virulence. Understanding the mechanisms of the antibacterial T6SS in *S. maltophilia* complex could lead to the development of novel treatment strategies against strains resistant to antibiotics.

## MATERIALS AND METHODS

### Bacterial strains and growth conditions

*S. maltophilia* complex STEN00241 and *P. aeruginosa* strain PSA01136 were previously isolated and sequenced as part of a bacterial whole genome sequencing surveillance study (60). *E. coli* DH5α*, B. cenocepacia* K56-2, *P. aeruginosa* PA14, and *S. aureus* JE2 were used for co-culture assays with STEN00241. *P. aeruginosa* PA32 (CFBR509_Pae_20170525_S_EBPa32) was isolated from the sputum of a CF patient (58). *E. coli* SM10 was used to conjugate plasmids into *S. maltophilia.* Strains were routinely grown in LB medium at 37*°*C unless otherwise indicated. The following antibiotic concentrations were used: chloramphenicol (20 μg/ml), imipenem (20 μg/ml or 32 μg/ml for imipenem monohydrate), gentamicin (75 μg/ml), and tetracycline (20 μg/ml). All strains used in this study are listed in Supp. Table 2.

### Construction of phylogenetic trees and ANI matrix

*S. maltophilia* protein sequences from both RefSeq and Genbank NCBI databases were retrieved on 09/04/2022 and used to create a local BLAST database. A local BlastP search was conducted using the TssC protein sequence of *X. citri.* Incomplete hits were eliminated from further analyses. *S. maltophilia* TssC sequences and TssC sequences from *X. citri, P. aeruginosa* PAO1 (TssC1 and TssC2), *Vibrio cholerae* V52, *Burkholderia cenocepacia* K56-2, *Acinetobacter baumannii* DSM 30011 were aligned using MUSCLE (71). Alignments were used to construct a phylogenetic tree in PhyML with a Bayesian Information Criterion Smart Model Selection (72, 73). Branch supports were calculated using the aLRT SH-like fast likelihood-based method (72). The final tree was created with iTOL (74). *S. maltophilia* genomes encoding a TssC protein were also confirmed to encode TssB and TssM T6SS proteins. The average nucleotide identity (ANI) of *S. maltophilia* complex genomes harboring T6SS genes was calculated using FastANI and a pairwise ANI matrix was built using ANIClustermap (75).

### Molecular biology

Standard molecular biology techniques were used to construct plasmids and PCR products. All restriction enzymes, DNA polymerases, and Gibson mixes were utilized according to manufacturer instructions. Plasmid constructs and PCR products were sequenced by GENEWIZ (Azenta Life Sciences, Chelmsford, MA, USA).

### Genetic manipulation of *S. maltophilia*

To engineer the *S. maltophilia* complex STEN00241 Δ*tssM* mutant, a similar allelic exchange method to the one described by Welker et al. was used (76). Briefly, 1000 bp upstream and downstream of the *tssM* gene (including the start and stop codons) were assembled on the pEX18Tc suicide vector using Gibson assembly. The pEX18Tc-Δ*tssM* construct was transformed into *E. coli* SM10 and then conjugated into *S. maltophilia* STEN00241. Conjugants were identified by growth on tetracycline and imipenem and then plated onto freshly made 15% sucrose LB plates (with no NaCl). Colonies were screened by PCR and confirmed by Sanger sequencing. Plasmids used in this study are listed in Supp. Table 2 and primers are listed in Supp. Table 3.

### Co-culture assays

Overnight cultures of the indicated strains were made from single colonies and were diluted 1:100 in fresh LB medium. Cultures were grown for 4 hours and set to an OD_600_ = 1.0. For *E. coli* DH5α carrying the tetracycline-resistant pEX18Tc plasmid, tetracycline was added to the growth media and cultures were washed with fresh LB before co-culturing with *S. maltophilia*. A 1x volume of target strains was mixed with a 10x volume *S. maltophilia,* centrifuged, and mixtures were resuspended in a 1x volume of LB. 5 µL of the mixed strains or of the indicated strains alone were spotted on LB plates. Co-cultures were then incubated at 37°C (or 25°C where indicated) for 20 hours. Colonies were excised, placed in 1 mL of LB, vortexed for 30 seconds, serially diluted, and plated on the following selective plates: tetracycline (20 μg/ml) to select for *E. coli,* chloramphenicol (20 μg/ml) to select for *P. aeruginosa*, gentamicin (75 μg/ml) to select for *B. cenocepacia*, *Staphylococcus* isolation agar (Trypticase soy agar with 7.5% NaCl) to select for *S. aureus,* and imipenem to select for *S. maltophilia*. For co-cultures in liquid media, 5 µL of co-cultures or strains alone were inoculated in 3 mL of liquid LB, serially diluted, and plated on selective plates. After overnight growth, the number of colonies was determined.

### Confocal microscopy

Plasmid pMRP9-1 (GFP+) (77) was introduced into *P. aeruginos*a PSA01136 and plasmid pMP7605 (mCherry+) (78) was introduced into *S. maltophilia* complex STEN00241 (both WT and T6SS-) by electroporation. Overnight cultures of WT and Δ*tssM S. maltophilia* complex STEN00241 expressing mCherry+ and *P. aeruginos*a PSA01136 expressing GFP+ were inoculated into LB medium from single colonies. Overnight cultures were diluted 1:100 in fresh LB medium, grown for 4 hours, washed with LB, and set to an OD_600_ = 1.0. A 1x volume of PSA01136 expressing GFP+ was mixed with a 10x volume STEN00241 mCherry+, centrifuged, and resuspended in 1x LB volume. 5 µL of the mixed strains or the indicated strains alone were spotted on dry LB plates and grown overnight. Colonies were visualized using a Zeiss LSM 710 upright microscope equipped with a Plan-Apochromat 10x objective. A 488 nm laser was used to observe *P. aeruginos*a PSA01136 expressing GFP+ and a 555 nm laser was used to observe *S. maltophilia* expressing mCherry+. Images were analyzed in FIJI. Representative images of three biological replicates are shown.

### Hcp immunoblotting

Polyclonal rabbit antibodies against the STEN00241 Hcp peptide N-LLQPRSATASTSGG-C were generated by Genescript. Overnight cultures of WT or Δ*tssM S. maltophilia* were washed with LB and back-diluted to an OD_600_ = 0.01. Strains were grown at 37°C to an OD_600_ ≈ 0.7. For cell fractions, 300 µL of cultures were centrifuged and cell pellets were resuspended in 100 µL of 1x Laemmli buffer. Samples were incubated at 99°C for 30 minutes. For supernatant fractions, cultures were centrifuged for 10 minutes at 2000 x g, supernatants were filter sterilized twice using 0.22 µM filters, and 700 µL of the filtered supernatants were incubated with trichloroacetic acid at a final concentration of 20% overnight at 4°C. Samples were then centrifuged for 10 minutes and washed three times with cold acetone. Precipitated proteins were resuspended in 150 µL of 1x Laemmli buffer in SDS buffer and incubated at 99°C for 60 minutes. Proteins from both cellular and supernatant fractions were separated on an SDS-PAGE gel and transferred to a PVDF membrane. The membrane was probed with primary antibodies against Hcp (1:1000) and RNA polymerase subunit alpha (1:2000), and secondary antibodies against rabbit (for Hcp) and mouse (for RNA polymerase subunit alpha) (1:5000, LI-COR Biosciences). The membrane was imaged using a Biorad ChemiDoc imager and analyzed in FIJI.

## ACKNOWLEDGMENTS

This work was supported by grants from the Cystic Fibrosis Foundation (GOLDBE19P0 to J.B.G, WHITEL20P0 to J.B.G, and CRISAN22F0 to C.V.C). This work was also supported by NIH grants R21-AI48847 to J.B.G and R01-AI27472 to D.V.T.). We thank the NIH IRACDA Fellowships in Research and Science Teaching (FIRST) program at Emory University for financial support to C.V.C.

We thank members of the Goldberg Lab, especially Rachel Done, Dr. Rebecca P. Duncan, Dr. Dina A. Moustafa, Justin Luu, Ashley Alexander, and Dr. Vishnu Raghuram for advice and assistance with experiments. We are grateful to the Emory University Integrated Cellular Imaging Core for help with confocal microscopy experiments. We also thank Dr. Lee H. Harrison for providing *S. maltophilia* and *P. aeruginosa* isolates collected from the EDS-HAT study.

